# Speak or shout? Nonverbal vocalizations ensure rapid detection of emotions in vocal communication

**DOI:** 10.1101/2025.06.18.660505

**Authors:** Marc D. Pell, Haining Cui, Yondu Mori, Xiaoming Jiang

**Author notes:** Corresponding author: Marc D. Pell, PhD, 2001 avenue McGill College, 8th floor, Montréal, Québec, H3A 1G1, Canada.

## Abstract

Human vocal expressions of emotion can be expressed nonverbally, through vocalizations such as shouts or laughter, or speakers can embed emotional meanings in language by modifying their tone of voice (“prosody”). Is there evidence that nonverbal expressions promote “better” (i.e., more accurate, faster) recognition of emotions than speech, and what is the impact of language experience? Our study investigated these questions using a cross-cultural gating paradigm, in which Chinese and Arab listeners (n=25/group) judged the emotion communicated by acoustic events that varied in duration (200 milliseconds to the full expression) and form (vocalizations or prosody expressed in listeners’ native, second or foreign language). Accuracy was higher for vocalizations overall, but listeners were markedly more *efficient* to form stable categorical representations of the speaker’s emotion from vocalizations (M = 417ms) than native prosody (M = 765ms). Language experience enhanced recognition of emotional prosody expressed by native/ingroup speakers for some listeners (Chinese) but not all (Arab), emphasizing the dynamic interplay of socio-cultural factors and stimulus quality on prosody recognition which occurs over a more sustained time window. Our data show that vocalizations are functionally suited to build robust, rapid impressions of a speaker’s emotion state unconstrained by the listener’s linguistic cultural background.

## Introduction

Human vocal expressions of emotion take two principal forms: they can be expressed nonverbally through laughter, shouts, or other emotionally relevant *vocalizations*; or speakers can embed emotional meaning in language by modifying suprasegmental acoustic features of their “tone of voice” (*speech prosody*) ^1^. Understanding what is shared and what is distinct about these two communication subsystems—in terms of their expressive form, social functions, neurocognitive underpinnings, and impact on listeners—has been the focus of recent work ^2–4^. Our study adds to this database by comparing emotion recognition from vocal expressions that vary in *exposure duration* and *form* (vocalizations vs. prosody expressed in a listener’s native, second or foreign language), allowing new insights about the nature and time course of processes that lead to emotion recognition in the vocal channel.

### One channel, two forms of expressions

Emotional vocalizations and emotional speech prosody share much in common; both consist of dynamic acoustic patterns that vary in perceived pitch, energy, quality, and temporal sound properties that progressively gain significance over time ^5^. There is evidence that both forms of expression exploit a common set of acoustic features that allow listeners to differentiate and attribute a range of discrete emotion states to the speaker ^4,6^. These mappings allow ‘basic’ emotions such as anger, fear, happiness and many other meanings to be successfully detected in the voice at levels far exceeding chance, both within and across cultural boundaries ^7–12^.

Still, it seems obvious that saying, *“I don’t know what to do”* in a sad voice, or beginning to sob uncontrollably, are quite different exemplars of “sadness” that will have distinct perceptual and interpersonal effects. At the production stage, vocalizations allow greater flexibility and acoustic variability in their expression than speech-embedded emotions, which are constrained by simultaneous demands on linguistic production ^13^. As a result, nonverbal signals are believed to encode emotions with greater perceptual clarity, promoting higher recognition rates than prosody when formally measured in experimental tasks ^2,4^. Vocalizations also appear to be processed with greater automaticity ^3,14,15^ and to capture preferential attention ^16,17^ over speech. The apparent primacy of emotional vocalizations in perception could reflect the fact that nonverbal signals emanate from a biologically primitive, reflexive call system adapted by humans (and other species) for survival, whereas emotions in speech are cognitively mediated and under significant voluntary control ^18–20^. As such, processes of socialization—including cultural preferences, language familiarity, and other forms of learning—are believed to exert stronger effects on how emotional prosody is recognized in the vocal channel than nonverbal expressions ^7,10,21^.

### On the time course of vocal emotion recognition

But how does the form of vocal expressions alter the *path* to emotion recognition over time, and in what way(s) do these operations depend on listener experience (e.g., previous exposure or familiarity with particular event types)? Surprisingly, direct comparisons of vocalizations and speech prosody are still few. One way to infer how and *when* vocal emotions are recognized is to measure event-related brain potentials (ERPs) evoked by a vocal stimulus. When listeners are presented prosody in their native language, there is evidence that the brain differentiates discrete emotional qualities of speech beginning 200 milliseconds post-onset of an utterance ^3,22^, at least in sufficient terms to facilitate deeper cognitive analysis of the event’s contextual meaning at later timepoints (see ^23^ for a recent discussion). ERP studies that have presented vocalizations tend to report even earlier cortical responses, with emotion-related changes in activity often beginning 100ms post-onset of vocalizations ^15,24^. These studies suggest that perceptual and cognitive operations for ‘recognizing’ emotion from voices proceed rapidly, but that vocalizations are differentiated at *an earlier time point* by the neurocognitive system than prosody. This claim is reinforced by an ERP experiment that compared the two expression types directly; vocalizations of anger, sadness, and happiness produced earlier and more qualitatively distinct brain responses in listeners than when these emotions were expressed prosodically in native-like pseudo-utterances ^3^.

Another way to illuminate the time course of vocal emotion recognition is by “gating” auditory stimuli, i.e., presenting time-limited excerpts of a vocal stimulus (e.g., the first 200 or 400 milliseconds), to gauge their effects on perception and behaviour. Using gated prosodic stimuli in a novel voice-face priming paradigm, Pell and colleagues ^25,26^ concluded that English listeners require at least 400ms of speech exposure to prime decisions about an emotionally-related face, suggesting that discrete meanings of prosody are activated and implicitly “recognized” from ∼400ms of acoustic information (see also ^27^). Interestingly, when emotions were expressed in a foreign language (Arabic), English listeners required prolonged exposure to the prosodic input (>600ms) for priming to occur ^28^. These findings suggest that language familiarity is a critical factor governing the recognition of speech-embedded emotions and alters its recognition time course ^29^.

Other experiments have probed these questions using an auditory gating paradigm adapted from ^30^, whereby participants judge vocal emotion expressions presented in time- or structure-based increments of increasing duration over the course of a study. For example, participants render forced-choice judgements about the meaning of an acoustic stimulus after hearing the first 100ms, 200ms, or 300ms of the same event. This approach allows researchers to estimate how much acoustic input listeners need to form stable categorical representations of emotions based on a specific stimulus duration or “gate”. Studies that have gated emotional prosody in languages such as English, Hindi, or Swedish show that speech-embedded emotions are accurately recognized at notably different latencies or “speeds” ^29,31–34^. While patterns vary from study to study due to differences in how gates were defined, most gating studies conclude that listeners need at least 400-500ms of emotional prosody for recognition to begin to stabilize (“emotion identification point”), meaning that participants can identify the target meaning at this timepoint and do not change their mind at longer exposures ^31,32,34^. The time course for recognizing specific emotions from prosody varies markedly: anger, sadness and (sometimes) fear tend to be isolated earliest from ∼500-800ms of acoustic information, whereas other emotions (e.g., happiness, disgust, interest) often require 1-2 seconds of speech cues to identify ^29,32,34^. Moreover, certain speech-embedded emotions, particularly happiness, seem to depend heavily on linguistic structure and can only be isolated when listeners integrate acoustic cues provided toward the end of an utterance ^7,33,35^.

Subsequent experiments that have gated vocalizations report that stable emotion representations can typically be formed after hearing ∼250-350ms of acoustic input, much earlier than for prosody ^36–38^. Castiajo & Pinheiro (2019) observed large increases in target hit rates for 10 different emotional vocalizations that lasted between 200-300ms, with fastest recognition of amusement (laughter) and slowest recognition of fear. However, this literature is again marked by many discrepancies owing to differences in the number and type of emotions studied and the way that auditory stimuli were gated for presentation (e.g., 33ms vs. 100ms intervals), among other methodological differences. To address these issues, our study undertook a direct test of how emotion recognition unfolds from vocalizations vs. speech prosody using the gating paradigm to determine whether nonverbal stimuli are recognized at an earlier timepoint than speech prosody, as suggested by recent neurophysiological findings ^3^.

As a secondary goal, we sought to shed light on how *familiarity* with a language influences the recognition of speech-embedded emotions. According to Dialect theory, socially-constructed forms of emotional communication, such as prosody, are shaped by cultural “styles” which provide listeners an advantage to recognize emotions expressed by native (‘ingroup’) speakers ^39^. In the only emotional gating study to implement a cross-cultural design, Jiang and colleagues ^29^ required groups of English Canadian and Indian listeners to recognize four emotions (anger, fear, happiness, sadness) from both English and Hindi pseudo-utterances gated to six exposure durations (200ms, 400ms, 500ms, 600ms, 700ms, full utterance). Results showed that recognition accuracy was significantly higher for native prosody in each group (ingroup advantage), emphasizing the importance of language experience on emotional prosody recognition ^40^. In addition, emotion identification points occurred earlier for native prosody, irrespective of whether the non-native language was considered foreign to the listener (Canadians judging Hindi) or the listener’s second language/L2 (Indians judging English). Interestingly, the Indian participants’ ability to recognize emotions in L2-English was positively associated with their English proficiency level, although this relationship is not consistently reported elsewhere in the literature (cf. ^41–43^. These findings motivate a deeper look at the impact of language familiarity—i.e., whether emotions are expressed in a listener’s native, second, or a completely foreign language—on both the accuracy and time course of emotional prosody recognition.

### Aim of the current study

Here, we investigated how emotions are recognized from vocalizations versus speech prosody using an adapted version of Jiang et al.’s ^29^ cross-cultural gating paradigm. Our new design simultaneously allowed us to evaluate the impact of language familiarity on recognition performance within and between two distinct groups: native speakers of Mandarin-Chinese and Arabic, all of whom were proficient second-language speakers of English. Based on our analysis above, we predicted that recognition accuracy would be higher and stabilize at earlier timepoints for vocalizations than for native prosodic expressions in each group; emotion-specific recognition trajectories in each condition were likely to vary, but these patterns should be more similar between groups when judging vocalizations than emotional prosody, which is shaped by linguistic and cultural variables to a greater extent ^40^. For speech-embedded emotions, it was expected that language familiarity would enhance accuracy and speed of emotional prosody recognition when each listener group judged their native language, consistent with the ingroup advantage (native > foreign and L2-English), but that high proficiency in L2-English could also enhance emotional prosody recognition in some manner over foreign prosody ^29,41,43^.

## Materials and Methods

### Participants

Fifty young adults judged all vocal expressions presented in the study; assuming medium effect sizes, this sample size achieves power exceeding .95 to detect differences in event type if mixed ANOVAs had been employed (this estimation may be considered the lower bound of actual power achieved in the study using linear mixed effects models (Kuznetsova et al., 2017). Participants were recruited from the greater Montréal region on the basis of speaking Arabic (n=25, 15F/10M, Mean Age = 21.5 ± 3.4, Mean Education = 15.6 years ± 3.8) or Mandarin-Chinese (n=25, 18F/7M, Mean Age = 22.2 ± 4.2, Mean Education = 15.6 years ± 2.4) as a native language, and for having high proficiency in English as a second language. Participants were students or recent immigrants who moved to Canada as adults and had lived in Montréal for less than five years (Median duration in Canada: Arab group = 18 months, range = 4-58 months; Chinese group = 12 months, range = 1-24 months). Most participants arrived in Canada between the ages of 19-21. Arab participants were born and raised in several Arabic-speaking countries (Syria, Jordan, Bahrain), although the majority (20/25) spoke variants of Levantine Arabic. Chinese participants were born and raised in different regions of mainland China (Shanghai, Beijing, Shenzhen). All participants learned English in school as a second language (L2-English) from an early age (Mean Age of English onset: Arab = 5.7 years ± 3.4; Chinese = 8.1 years ± 3.3). L2-English proficiency was characterized through a series of self-report measures gathered at the onset of the study; all participants in each group rated their ability to speak and listen in English as high (group means ranged between 7.7-9.5 on a 10-point proficiency scale, see Table 1). Many participants in each group knew additional languages (e.g., French, Farsi, Cantonese). To enter the study, it was verified that no participant had any knowledge of the language designated as “foreign” for that group in the experiment (i.e., Arabic for Chinese participants; Chinese for Arab participants). All participants reported normal hearing. Recruitment began on 07/21/2016 and ended on 05/26/2017. Voluntary written consent was obtained prior to the study, which was approved by the Faculty of Medicine and Health Sciences Institutional Review Board, McGill University.

**Table 1.** Demographic characteristics of participants in the Chinese and Arab groups.

| *Variable* | **Chinese Group**<br>**(n=25)** |  | **Arab Group**<br>**(n=25)** |  |
| --- | --- | --- | --- | --- |
|  | *Mean* | *SD* | *Mean* | *SD* |
| Sex (f/m) | 18/7 | -- | 15/10 | -- |
| Age at testing (years) | 22.2 | 4.2 | 21.5 | 3.4 |
| Age of arrival to Canada (years) | 21.2 | 8.1 | 19.8 | 4.2 |
| Time in Canada (months) | 12.4 | 4.6 | 21.8 | 16.2 |
| Education (years) | 15.0 | 2.4 | 15.6 | 3.8 |
| Age of L2-English onset (years) | 8.1 | 3.3 | 5.8 | 3.4 |
| Self-reported L2-English Proficiency<br>(scale 1-10) |  |  |  |  |
| Listening | 8.6 | 1.3 | 9.5 | 0.8 |
| Speaking | 7.7 | 1.5 | 8.8 | 1.6 |
| Reading | 8.2 | 1.4 | 9.4 | 0.9 |
| Writing | 7.2 | 1.9 | 8.8 | 1.2 |

### Materials

The stimuli were digital auditory recordings of vocally expressed emotion lasting ∼1-3 seconds in duration, produced by a variety of speakers. Materials were divided into two main types of vocal events: nonverbal vocalizations and speech-embedded emotional expressions (henceforth referred to as *vocalization* and *speech prosody*). Speech prosody was further defined by its language of expression (Arabic, Mandarin, English) and the contextual relationship or familiarity of each language to the perceiver (native, L2-English, foreign). Vocalizations and prosody expressed in each language communicated one of four emotions (anger, fear, sadness, happiness); recordings were taken from established inventories used actively in the literature (see below). All recordings were elicited in laboratory settings using lay speakers/actors and then validated to establish the perceptual reliability of emotional meanings encoded by particular items according to objectives of the original authors. Perceptual data from the original studies were used here to select a diverse but controlled set of exemplars suitable for comparing the effects of vocal event type on emotion recognition using the auditory gating paradigm.

Vocalizations – Nonverbal vocalizations took the form of growls or shouts (anger), cries (fear), sobbing or wailing (sadness), and laughter or contentment sounds (happiness). All stimuli were nonverbal in nature, although some resembled sustained vowels or nasal sounds found in most languages (“aah” or “mmm”). Items were selected from three separate corpora ^21,44,45^ to increase generalizability of results and allow finer selection of items to match features of the speech stimuli (vocalizations were produced by French Canadian, European Portuguese, and British English speakers). Items were selected for achieving high consensus about the target emotion according to the original study design (accuracy for all stimuli > 5 times chance). Items which were too brief for effective gating or much longer than the duration of the speech stimuli were excluded (Median duration of vocalizations = 1448ms, range = 715-2376ms). For vocalizations representing “happiness”, emotional meaning labels validated in the original inventories often differed (e.g., “pleasure”, “contentment”, “amusement”, “happiness”). As these terms likely refer to a range of positive emotions that are communicated nonverbally ^46^, we included two subtypes of “happy” vocalizations to compare with speech prosody: happiness-*amusement*, which was characterized uniquely by laughter sounds; and happiness-*pleasure,* which was composed of vocal sounds of pleasure or contentment (e.g., “mmm”, “aah”). Ten items were selected for anger, fear, and sadness and 20 items for happiness (10 amusement, 10 pleasure). This totalled 50 vocalizations, produced by a variety of speakers (7-10 speakers/emotion, half female, half male, 18 distinct voices in total).

Speech prosody *–* Stimuli in the speech condition were emotionally-inflected pseudo-utterances composed of 7-11 syllables/characters produced by native speakers of Arabic, Mandarin-Chinese, or English ^12,47^. Pseudo-utterances, which mimic linguistic properties of a language while restricting emotion to the speaker’s prosody, have been used in previous gating studies ^29,31,32^ and broadly in the prosody literature (see ^5^ for an overview). Pseudo-speech stimuli in each of the three language conditions were constructed, recorded, and perceptually validated using virtually identical procedures, but involving native speakers-listeners of only the target language (see ^47^ for details). For each recorded language, we selected expressions of one female and one male speaker producing six different utterances to convey each emotional target (anger, fear, sadness, happiness, 12 items/emotion), yielding 48 tokens/ language (2 speakers x 4 emotions x 6 utterances). Items were again selected for having high emotion recognition rates when judged by a group of native listeners in the original study (minimum 3x chance accuracy based on a seven forced-choice task). In addition, items were chosen to mitigate gross differences in native emotion recognition accuracy across language sets to the extent possible (Mean emotional target recognition by native listeners: Range = 55-74% for Arabic, 64-82% for Mandarin, 64-80% for English). The selected speech stimuli varied naturally in duration according to the emotion expressed and characteristics of each language but were roughly similar in overall duration to items in the vocalization condition (Median duration = 1493ms, range = 834-2900ms). A total of 144 utterances (3 languages x 48 items) were selected in the speech prosody condition for gating.

### Experimental design and procedures

Gate construction *–* Vocalizations (n=50) and speech prosody (n=144) were edited using Praat speech analysis software to standardize the peak volume of all sound files (75dB) and to segment each stimulus into four additional gates which varied in duration. Each item was cut from its acoustic onset to isolate the initial 200ms, 400ms, 500ms, and 600ms of the stimulus, which were saved as separate .wav files and manually edited to eliminate any noise artefacts (clipping noise). This procedure yielded five gate conditions per stimulus, the final gate always being the original unedited sound; these gates are referred to as G200, G400, G500, G600, and GFull. The choice of gate durations was informed by emotion identification points reported in previous gating studies ^32,36^ as well as Jiang et al.’s ^29^ cross-cultural study which reported effects of language familiarity on prosody recognition in the 400-600ms latency range. The gating process created 250 distinct trials for presentation in the vocalization condition (50 items x 5 gates) and 720 trials in the speech condition (3 languages x 48 items x 5 gates).

Testing procedures *–* Participants were tested in a quiet laboratory, individually or in small groups (2-4 people), seated at individual workstations. Instructions about the experiment were first provided verbally in English. Participants were told that they would hear vocal sounds or utterances that could sound familiar but would not make sense; they were instructed to pay attention to the speaker’s voice to decide what emotion the sound conveys *and* how confident they felt about their decision. Participants were told that stimuli would begin short and would sound “cut off”, and that sounds would increase in duration over the experiment. They were told to choose the label that *best fits* their impression of the emotion being expressed or to guess when they were unsure. Once participants had begun the study, written instructions and all other features of the experiment (e.g., emotion labels) were only presented in the participant’s native language (Arabic or Mandarin).

Listeners heard stimuli over volume adjustable headphones controlled by Superlab 5.0 presentation software (Cedrus, CA). Each trial began with an inter-stimulus interval (ISI) of 1500ms, a 500ms fixation cross, another 1500ms ISI, the auditory target, followed by a response screen. Participants clicked one of five emotion labels (anger, happiness, fear, sadness, neutral) displayed in the participant’s native language (for Mandarin: 高兴, 生气, 害怕, 难过, 中性; for Arabic, ﺳﻌﺎﺩﺓ,ﻏﻀﺐ,ﺧﻮﻑ,ﺣﺰﻥ,ﻣﺤﺎﻳﺪ.). Immediately after, participants saw a new screen and rated their confidence in their judgment along a 7-point scale (1= *not at all* to 7 = *very much*). Trials were presented in five blocks of increasing gate duration, starting with the shortest gate (G200) and ending with the full stimulus (GFull, ^30,32^. Within each gated block (e.g., G200), vocal event type (speech vs. vocalization) was blocked separately and counterbalanced for presentation order across participants (i.e., speech and vocalizations were each randomized within a gated block but were not intermixed). Prosodic expressions in the three language contexts (Arabic, Mandarin, English) were randomly intermixed within each gated block of the speech condition. Before judging stimuli at a particular gate, participants completed five practice trials; breaks were programed at the middle and end of each block. The experiment took 2.5 hours to complete, and participants received $25 CAD as compensation.

### Statistical analysis

Accuracy and latency of emotion recognition served as the two dependent measures of interest. Accuracy was estimated using Hu scores ^48^, the proportion of correct target responses assigned to each emotion adjusted for the number of items in each category and individual biases in category usage. Individual Hu scores were calculated for each emotion at each gate interval, separately for vocalizations and speech prosody judged in each language context. Recognition latency was estimated by calculating the Emotion Identification Point (EIP) for each item ^32^. The EIP is the gate at which a participant correctly recognized the target meaning of a stimulus without changing their response at longer exposures of the same event, expressed in milliseconds (200ms, 400ms, 500ms, 600ms or the actual full event duration, ranging from 834-2900ms). Thus, EIPs considered a maximum of 250 datapoints per emotion/group for vocalizations and 300 judgements per emotion/group for each speech prosody condition. Items that did not lead to stable recognition by GFull were scored as errors and excluded from EIP calculations ^32^. On average, 1642 observations (range = 1433-1838) contributed to the calculation of EIPs for each event type.

Linear mixed-effects models (LMM) were built to separately infer how accuracy (Hu score) and latency (EIP) measures were influenced by our experimental manipulations. Analyses were performed in R Studio (Version 4.2.2; http://cran.r-project.org) using the lme4 package ^49^ with Satterthwaite approximations method for providing degrees of freedom and F-statistics implemented in the *lmerTest* package ^50^. Most LMMs included some combination of the fixed factors: perceiver Group (Chinese, Arab), vocal Event type (vocalization, native, L2-English, foreign), Emotion (anger, fear, happiness, sadness), and/or Gate duration (G200, G400, G500, G600, GFull), with Participant entered as a random factor. Emotion was also entered as a random effect in later models, along with variability by participant to allow for a focus on the broad role of vocal event type and language familiarity. Given that our stimuli (both vocalizations and speech) varied considerably in length, GFull duration (in milliseconds) was included as a covariate in models considering the EIP data to eliminate any potential effect of stimulus duration on the latency data. The *emmeans* package was used for all pairwise post hoc comparisons.

## Results

Table 2 presents the unbiased accuracy rates (mean Hu score) at each stimulus duration, separately for the Chinese and Arab participants by event type and emotion. Table 3 shows the frequency/percentage of EIPs occurring at each stimulus duration across conditions, trials scored as errors (failure to recognize the target), and the mean EIPs for each emotion expressed in milliseconds (ms). The proportion of “neutral” responses assigned in each condition and the mean confidence ratings of each participant group when judging different forms of stimuli are furnished in Tables S1 and S2. The datasets generated during and/or analysed for the current study are available in the Open Science Framework repository (https://osf.io/43udq/?view_only=d2b4a9cdbb9541f3b292f2a8c490616d).

**Table 2.**
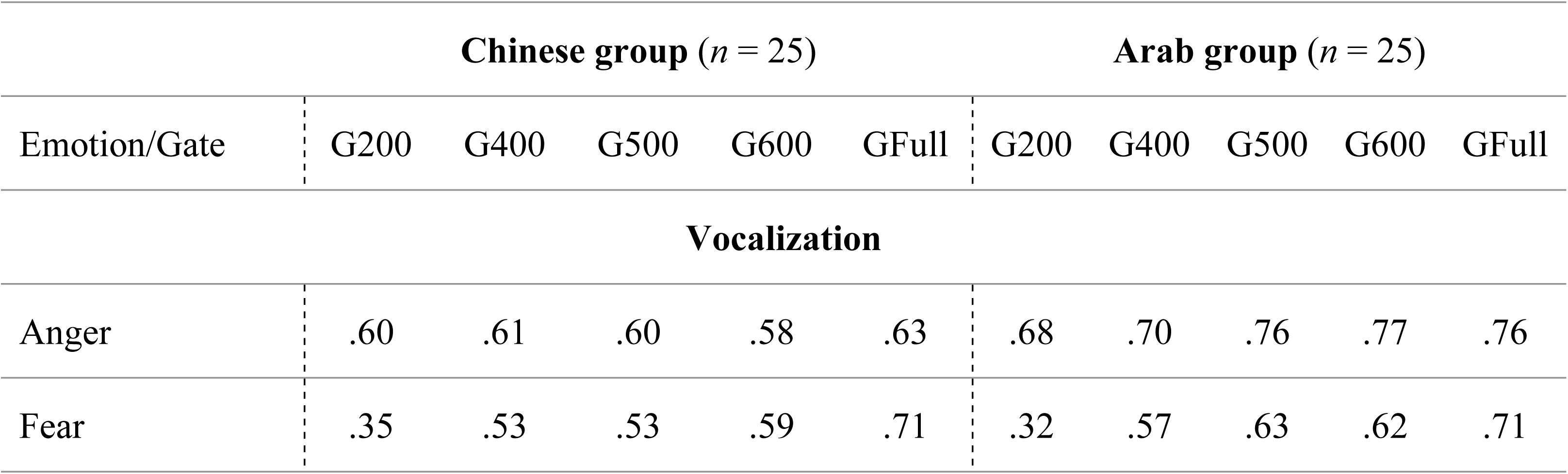
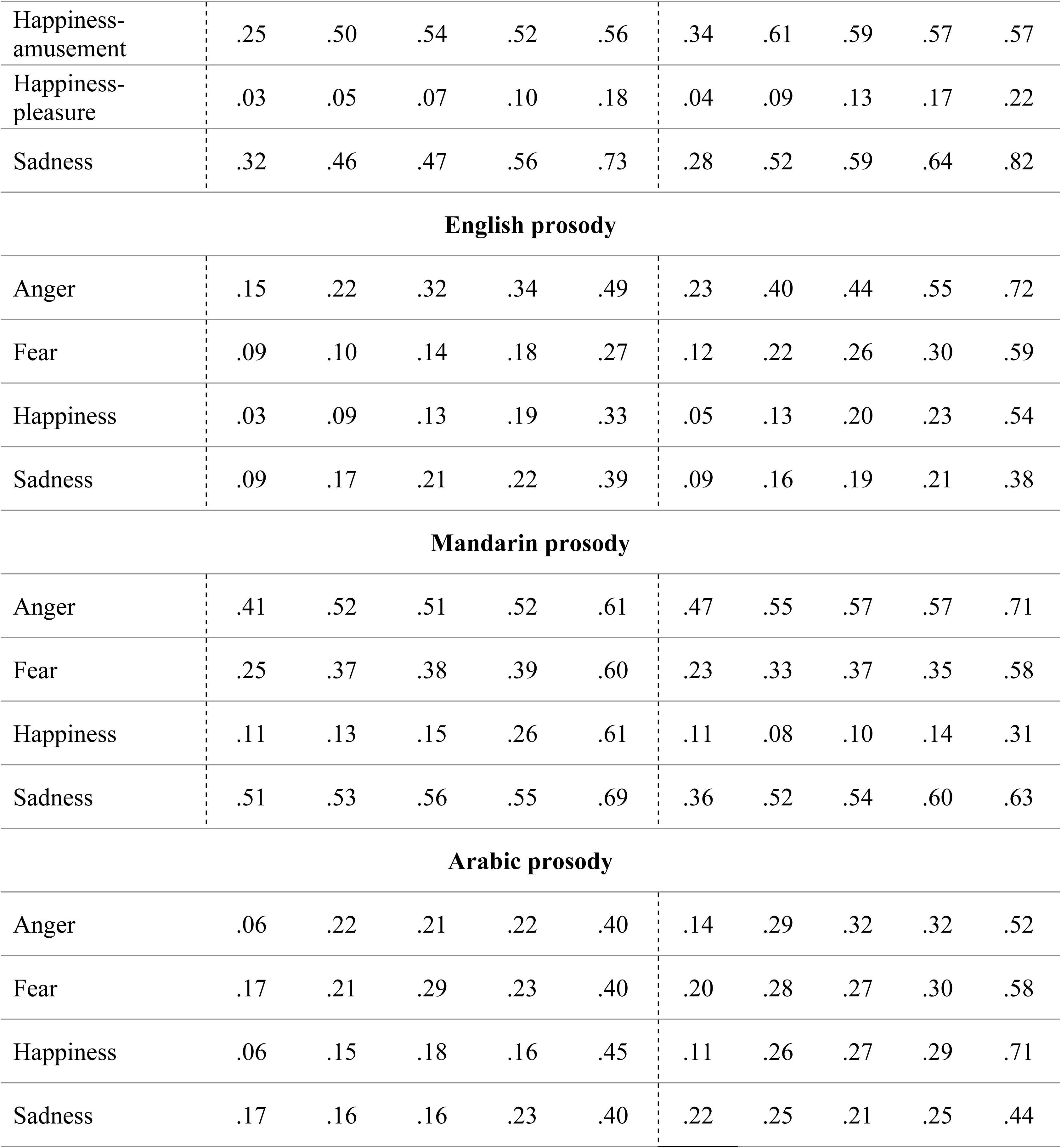
Unbiased hit rates (mean Hu scores) for Chinese and Arab participants by vocal event type, emotion, and gate duration.

| Emotion/Gate | Chinese group (*n* = 25) |  |  |  |  | Arab group (*n* = 25) |  |  |  |  |
| --- | --- | --- | --- | --- | --- | --- | --- | --- | --- | --- |
|  | G200 | G400 | G500 | G600 | GFull | G200 | G400 | G500 | G600 | GFull |
| **Vocalization** |  |  |  |  |  |  |  |  |  |  |
| Anger | .60 | .61 | .60 | .58 | .63 | .68 | .70 | .76 | .77 | .76 |
| Fear | .35 | .53 | .53 | .59 | .71 | .32 | .57 | .63 | .62 | .71 |
| Happiness-amusement | .25 | .50 | .54 | .52 | .56 | .34 | .61 | .59 | .57 | .57 |
| Happiness-pleasure | .03 | .05 | .07 | .10 | .18 | .04 | .09 | .13 | .17 | .22 |
| Sadness | .32 | .46 | .47 | .56 | .73 | .28 | .52 | .59 | .64 | .82 |
| **English prosody** |  |  |  |  |  |  |  |  |  |  |
| Anger | .15 | .22 | .32 | .34 | .49 | .23 | .40 | .44 | .55 | .72 |
| Fear | .09 | .10 | .14 | .18 | .27 | .12 | .22 | .26 | .30 | .59 |
| Happiness | .03 | .09 | .13 | .19 | .33 | .05 | .13 | .20 | .23 | .54 |
| Sadness | .09 | .17 | .21 | .22 | .39 | .09 | .16 | .19 | .21 | .38 |
| **Mandarin prosody** |  |  |  |  |  |  |  |  |  |  |
| Anger | .41 | .52 | .51 | .52 | .61 | .47 | .55 | .57 | .57 | .71 |
| Fear | .25 | .37 | .38 | .39 | .60 | .23 | .33 | .37 | .35 | .58 |
| Happiness | .11 | .13 | .15 | .26 | .61 | .11 | .08 | .10 | .14 | .31 |
| Sadness | .51 | .53 | .56 | .55 | .69 | .36 | .52 | .54 | .60 | .63 |
| **Arabic prosody** |  |  |  |  |  |  |  |  |  |  |
| Anger | .06 | .22 | .21 | .22 | .40 | .14 | .29 | .32 | .32 | .52 |
| Fear | .17 | .21 | .29 | .23 | .40 | .20 | .28 | .27 | .30 | .58 |
| Happiness | .06 | .15 | .18 | .16 | .45 | .11 | .26 | .27 | .29 | .71 |
| Sadness | .17 | .16 | .16 | .23 | .40 | .22 | .25 | .21 | .25 | .44 |

**Table 3.**
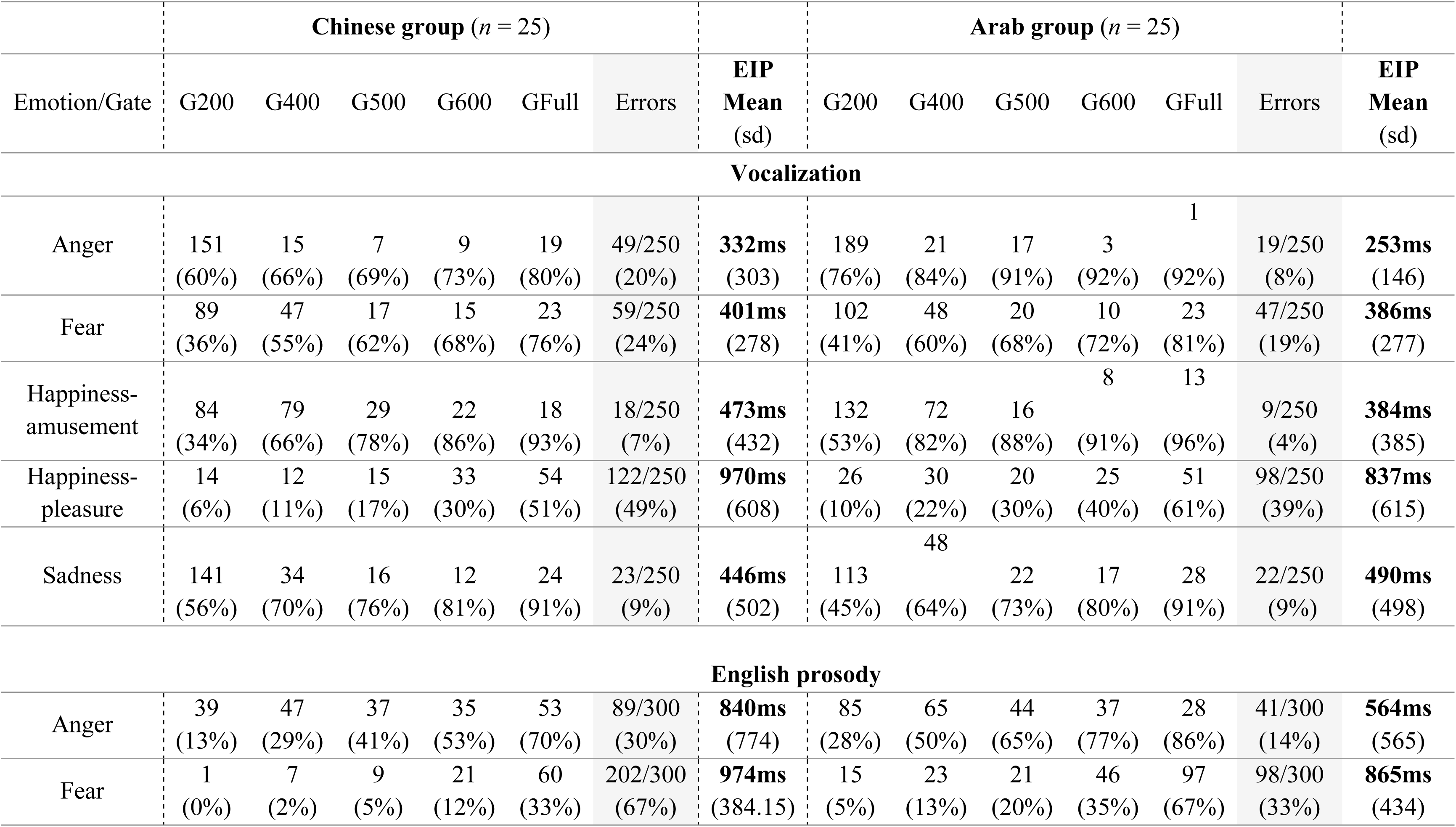
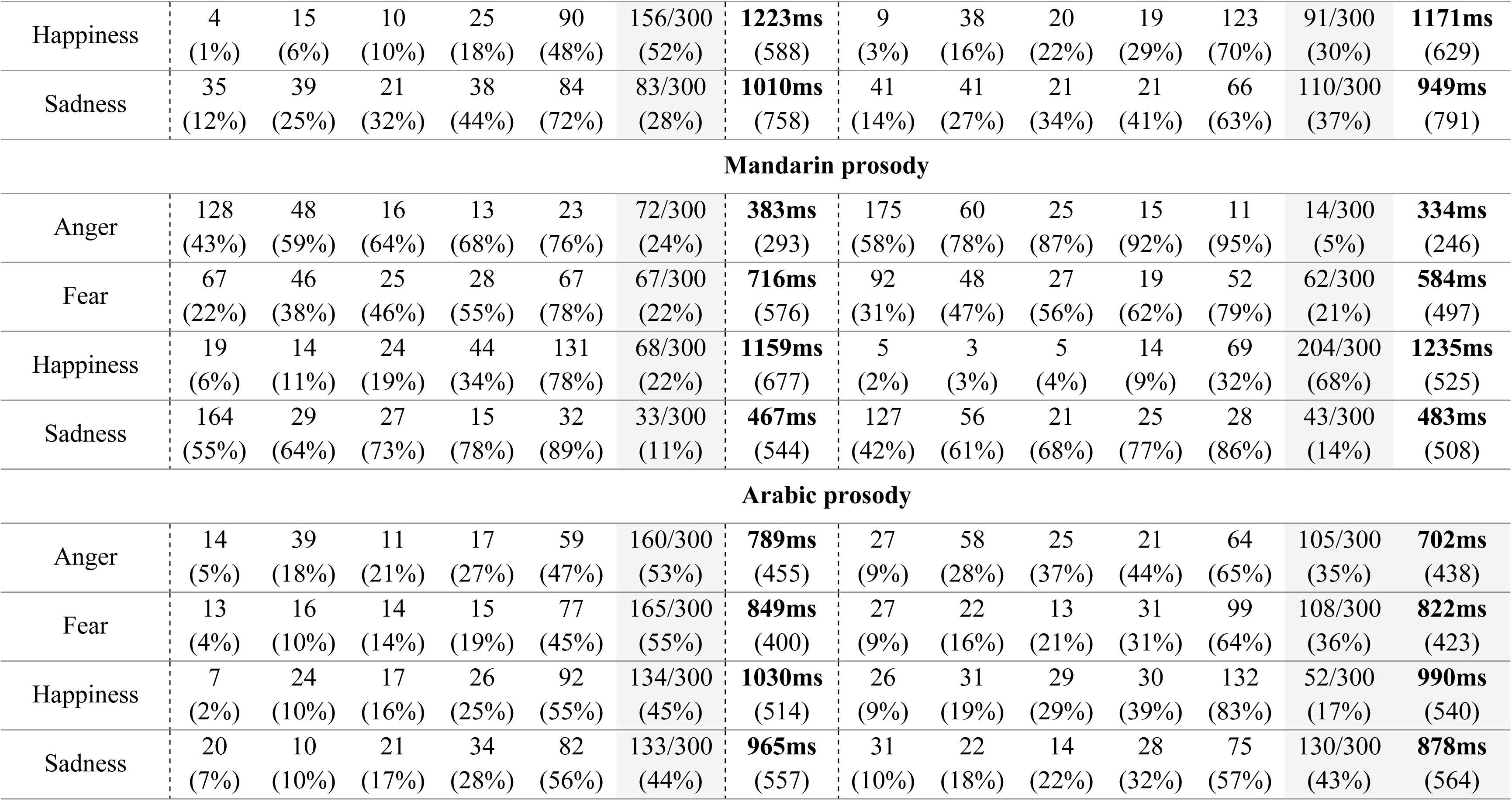
Frequency (+ cumulative percent) of stimuli recognized at each gate which contributed to the calculation of Emotion Identification Points (EIP), by group, event type and emotion. Shaded cells show the frequency (percent) of stimuli coded as errors (i.e., the emotional target was not identified by GFull for a given participant/stimulus). Mean EIPs (+ standard deviation) are presented in bold. EIPs considered a total of 250 observations/emotion (25 participants x 10 items) for vocalizations and 300 observations/emotion (25 participants x 12 items) for each prosody condition per group.

|  | Chinese group (n = 25) |  |  |  |  |  |  | Arab group (n = 25) |  |  |  |  |  |  |
| --- | --- | --- | --- | --- | --- | --- | --- | --- | --- | --- | --- | --- | --- | --- |
| Emotion/Gate | G200 | G400 | G500 | G600 | GFull | Errors | EIP<br>Mean<br>(sd) | G200 | G400 | G500 | G600 | GFull | Errors | EIP<br>Mean<br>(sd) |
| Vocalization |  |  |  |  |  |  |  |  |  |  |  |  |  |  |
| Anger | 151<br>(60%) | 15<br>(66%) | 7<br>(69%) | 9<br>(73%) | 19<br>(80%) | 49/250<br>(20%) | **332ms**<br>(303) | 189<br>(76%) | 21<br>(84%) | 17<br>(91%) | 3<br>(92%) | 1<br>(92%) | 19/250<br>(8%) | **253ms**<br>(146) |
| Fear | 89<br>(36%) | 47<br>(55%) | 17<br>(62%) | 15<br>(68%) | 23<br>(76%) | 59/250<br>(24%) | **401ms**<br>(278) | 102<br>(41%) | 48<br>(60%) | 20<br>(68%) | 10<br>(72%) | 23<br>(81%) | 47/250<br>(19%) | **386ms**<br>(277) |
| Happiness-amusement | 84<br>(34%) | 79<br>(66%) | 29<br>(78%) | 22<br>(86%) | 18<br>(93%) | 18/250<br>(7%) | **473ms**<br>(432) | 132<br>(53%) | 72<br>(82%) | 16<br>(88%) | 8<br>(91%) | 13<br>(96%) | 9/250<br>(4%) | **384ms**<br>(385) |
| Happiness-pleasure | 14<br>(6%) | 12<br>(11%) | 15<br>(17%) | 33<br>(30%) | 54<br>(51%) | 122/250<br>(49%) | **970ms**<br>(608) | 26<br>(10%) | 30<br>(22%) | 20<br>(30%) | 25<br>(40%) | 51<br>(61%) | 98/250<br>(39%) | **837ms**<br>(615) |
| Sadness | 141<br>(56%) | 34<br>(70%) | 16<br>(76%) | 12<br>(81%) | 24<br>(91%) | 23/250<br>(9%) | **446ms**<br>(502) | 113<br>(45%) | 48<br>(64%) | 22<br>(73%) | 17<br>(80%) | 28<br>(91%) | 22/250<br>(9%) | **490ms**<br>(498) |
| English prosody |  |  |  |  |  |  |  |  |  |  |  |  |  |  |
| Anger | 39<br>(13%) | 47<br>(29%) | 37<br>(41%) | 35<br>(53%) | 53<br>(70%) | 89/300<br>(30%) | **840ms**<br>(774) | 85<br>(28%) | 65<br>(50%) | 44<br>(65%) | 37<br>(77%) | 28<br>(86%) | 41/300<br>(14%) | **564ms**<br>(565) |
| Fear | 1<br>(0%) | 7<br>(2%) | 9<br>(5%) | 21<br>(12%) | 60<br>(33%) | 202/300<br>(67%) | **974ms**<br>(384.15) | 15<br>(5%) | 23<br>(13%) | 21<br>(20%) | 46<br>(35%) | 97<br>(67%) | 98/300<br>(33%) | **865ms**<br>(434) |
| Happiness | 4<br>(1%) | 15<br>(6%) | 10<br>(10%) | 25<br>(18%) | 90<br>(48%) | 156/300<br>(52%) | **1223ms**<br>(588) | 9<br>(3%) | 38<br>(16%) | 20<br>(22%) | 19<br>(29%) | 123<br>(70%) | 91/300<br>(30%) | **1171ms**<br>(629) |
| Sadness | 35<br>(12%) | 39<br>(25%) | 21<br>(32%) | 38<br>(44%) | 84<br>(72%) | 83/300<br>(28%) | **1010ms**<br>(758) | 41<br>(14%) | 41<br>(27%) | 21<br>(34%) | 21<br>(41%) | 66<br>(63%) | 110/300<br>(37%) | **949ms**<br>(791) |
| Mandarin prosody |  |  |  |  |  |  |  |  |  |  |  |  |  |  |
| Anger | 128<br>(43%) | 48<br>(59%) | 16<br>(64%) | 13<br>(68%) | 23<br>(76%) | 72/300<br>(24%) | **383ms**<br>(293) | 175<br>(58%) | 60<br>(78%) | 25<br>(87%) | 15<br>(92%) | 11<br>(95%) | 14/300<br>(5%) | **334ms**<br>(246) |
| Fear | 67<br>(22%) | 46<br>(38%) | 25<br>(46%) | 28<br>(55%) | 67<br>(78%) | 67/300<br>(22%) | **716ms**<br>(576) | 92<br>(31%) | 48<br>(47%) | 27<br>(56%) | 19<br>(62%) | 52<br>(79%) | 62/300<br>(21%) | **584ms**<br>(497) |
| Happiness | 19<br>(6%) | 14<br>(11%) | 24<br>(19%) | 44<br>(34%) | 131<br>(78%) | 68/300<br>(22%) | **1159ms**<br>(677) | 5<br>(2%) | 3<br>(3%) | 5<br>(4%) | 14<br>(9%) | 69<br>(32%) | 204/300<br>(68%) | **1235ms**<br>(525) |
| Sadness | 164<br>(55%) | 29<br>(64%) | 27<br>(73%) | 15<br>(78%) | 32<br>(89%) | 33/300<br>(11%) | **467ms**<br>(544) | 127<br>(42%) | 56<br>(61%) | 21<br>(68%) | 25<br>(77%) | 28<br>(86%) | 43/300<br>(14%) | **483ms**<br>(508) |
| Arabic prosody |  |  |  |  |  |  |  |  |  |  |  |  |  |  |
| Anger | 14<br>(5%) | 39<br>(18%) | 11<br>(21%) | 17<br>(27%) | 59<br>(47%) | 160/300<br>(53%) | **789ms**<br>(455) | 27<br>(9%) | 58<br>(28%) | 25<br>(37%) | 21<br>(44%) | 64<br>(65%) | 105/300<br>(35%) | **702ms**<br>(438) |
| Fear | 13<br>(4%) | 16<br>(10%) | 14<br>(14%) | 15<br>(19%) | 77<br>(45%) | 165/300<br>(55%) | **849ms**<br>(400) | 27<br>(9%) | 22<br>(16%) | 13<br>(21%) | 31<br>(31%) | 99<br>(64%) | 108/300<br>(36%) | **822ms**<br>(423) |
| Happiness | 7<br>(2%) | 24<br>(10%) | 17<br>(16%) | 26<br>(25%) | 92<br>(55%) | 134/300<br>(45%) | **1030ms**<br>(514) | 26<br>(9%) | 31<br>(19%) | 29<br>(29%) | 30<br>(39%) | 132<br>(83%) | 52/300<br>(17%) | **990ms**<br>(540) |
| Sadness | 20<br>(7%) | 10<br>(10%) | 21<br>(17%) | 34<br>(28%) | 82<br>(56%) | 133/300<br>(44%) | **965ms**<br>(557) | 31<br>(10%) | 22<br>(18%) | 14<br>(22%) | 28<br>(32%) | 75<br>(57%) | 130/300<br>(43%) | **878ms**<br>(564) |

Results were broken down as follows: we first characterized how accurately each group recognized specific emotions over time (i.e., as stimuli incrementally increased in duration from gate-to-gate), independently for vocalizations and when listeners judged emotional prosody in their native language. At the same time, we considered the latency associated with stable recognition of emotional targets for each event type, as inferred from the EIPs. These analyses allow comparisons with existing literature that describe the nature and time course of emotion recognition when vocalizations and speech prosody were studied separately. At a second stage, we directly tested whether listeners in our two groups displayed an overall *advantage* in accuracy and/or speed for vocalizations over native prosody, collapsing across emotion types to highlight the broad trends. At a final stage, we focused strictly on how emotions are recognized from prosodic stimuli to evaluate whether linguistic *familiarity* influences recognition of speech-embedded emotions in each group, comparing the performance measures in participants’ native language, second language (L2-English), and a foreign language (again collapsed across emotion types). Full statistical details of the LMMs and post hoc tests conducted on all significant main and interactive effects are reported in Tables S3 to S6. For expository purposes, only the *F*-statistics for significant effects are reported in the text.

### How does emotion recognition unfold from vocalizations?

#### Accuracy

Separate group analysis first looked at recognition accuracy for vocalizations (LMM: *HuScore (Vocalization) ∼ Emotion + Gate + Emotion * Gate + (1| Subject)*). Accuracy of each group depended on Emotion type (Arab: *F* = 287.81; *df* = 4, *p* < 0.001; Chinese: *F* = 275. 61, *df* =4, *p* < 0.001), Gate duration (Arab: *F* = 64.46, *df* = 4, *p* < 0.001; Chinese: *F* = 50.52, *df* =4, *p* < 0.001), and Emotion x Gate duration (Arab: *F =* 6.43, *df* = 16, *p* < 0.001; Chinese: *F* = 5.78, *df* =16, *p* < 0.001; S3 Table a-b). Fig. 1a-b shows that while emotional vocalizations had distinct recognition trajectories over time, the accuracy of Chinese and Arab listeners was similar in qualitative and quantitative terms. When accuracy at successive gate durations was compared, data show that anger (growls) was recognized at high levels based on 200ms of acoustic information (G200) and improved minimally as exposure increased. Happiness-amusement improved when stimulus duration increased from 200-400ms (*p*s < .001), without further improvements between gates to the full stimulus. Fear (screams) and sadness (sobs) also improved significantly between 200-400ms, and then again between G400 to the end of the stimulus (all *p*s < .001), suggesting a more incremental buildup of recognition for these signals. Happiness-pleasure was identified very poorly by both groups, and while accuracy improved between 200ms and the end of the utterance (*p*s < .002), recognition remained below chance performance levels for this emotion even when listeners heard the full stimulus.

**Figure 1.**
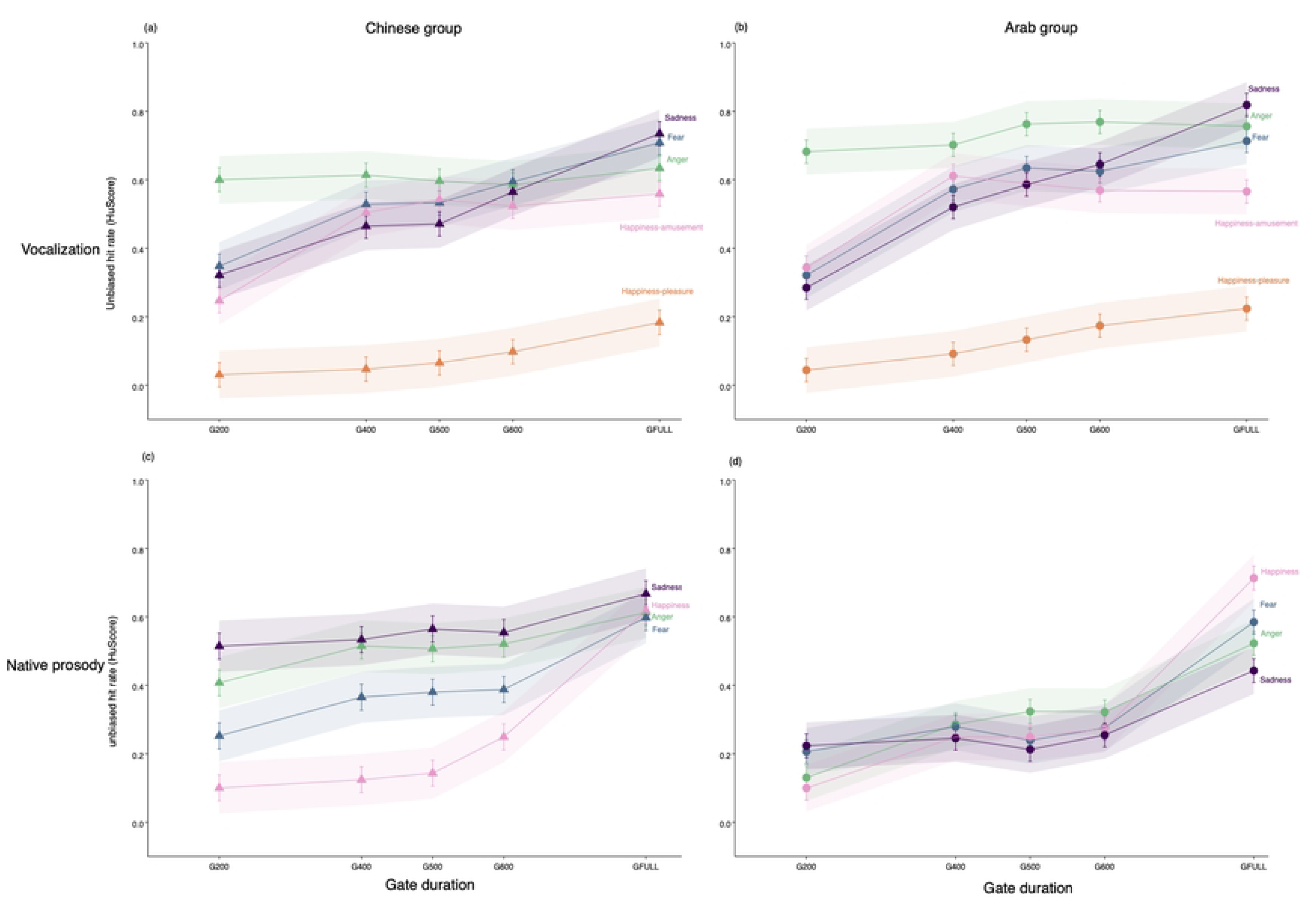
Recognition accuracy (HuScore) by Event type, Emotion, and Gate. Accuracy of each listener group to recognize emotions by vocal event type and stimulus (gate) duration. The top panels show the unbiased hit rates (Hu Scores) when nonverbal vocalizations were presented and the bottom panels showing corresponding data when emotional prosody was presented in the listener’s native language (Mandarin-Chinese or Arabic). The performance of Chinese listeners is shown on the left and Arab listeners is shown on the right.

Limiting stimuli to 200ms yielded superior recognition of anger versus all other vocalizations. Based on 400ms excerpts, anger, fear, and happiness-amusement were recognized at roughly similar rates by each group, exceeding sadness and happiness-pleasure. Virtually no changes occurred in the 400-500ms time range for either group. When stimuli lasted 600ms, only happiness-pleasure was recognized at inferior levels (Arab participants displayed a somewhat prolonged “anger” detection advantage over some vocalizations in the 200-600ms time range, Fig. 1b). Recognition of ungated vocalizations (GFull) was highest for anger, fear, and sadness, followed by happiness-amusement, with markedly inferior detection of happiness-pleasure. The fact that pleasure/contentment sounds, in contrast to laughter, were not reliably identified as “happiness” by either group irrespective of exposure time suggests an artefact of our forced-choice design for this category (S1 Table shows that these expressions were labelled as “neutral” approximately half of the time). For this reason, happiness-pleasure was not considered in further analyses; “happiness” in all subsequent models referred solely to amusement/ laughter sounds.

#### Latency

Analysis of EIPs (omitting happiness-pleasure) considered the time needed to form a stable representation of the four vocalizations between groups, controlling for differences in individual event duration (LMM: *EIPtime (Vocalization) ∼ Group + Emotion + Group* Emotion + GFullDuration + (1 | Subject)*); S3 Table c). EIPs varied by Emotion (*F* = 9.16, *df* = 3, *p* < 0.001) and Group x Emotion (*F* = 4.04, df = 3, *p* = 0.007). Fig. 2 shows that all vocalizations were reliably isolated by listeners in both groups within a narrow ∼300-500ms time window. Overall, fear (M = 500ms) required significantly more acoustic exposure to isolate than happiness-amusement (M = 355ms, *p <*001), anger (M = 394ms, *p*<.001), and sadness (M = 407ms, *p* =.019). The interaction was explained by small group differences in the significance of emotion-specific contrasts, and evidence that Arab listeners required less time to detect happiness (laughter) than the Chinese (*p =* .041, Fig. 2). There was no main effect of Group on recognition latencies for vocalizations (p = 0.25). .

**Figure 2.**
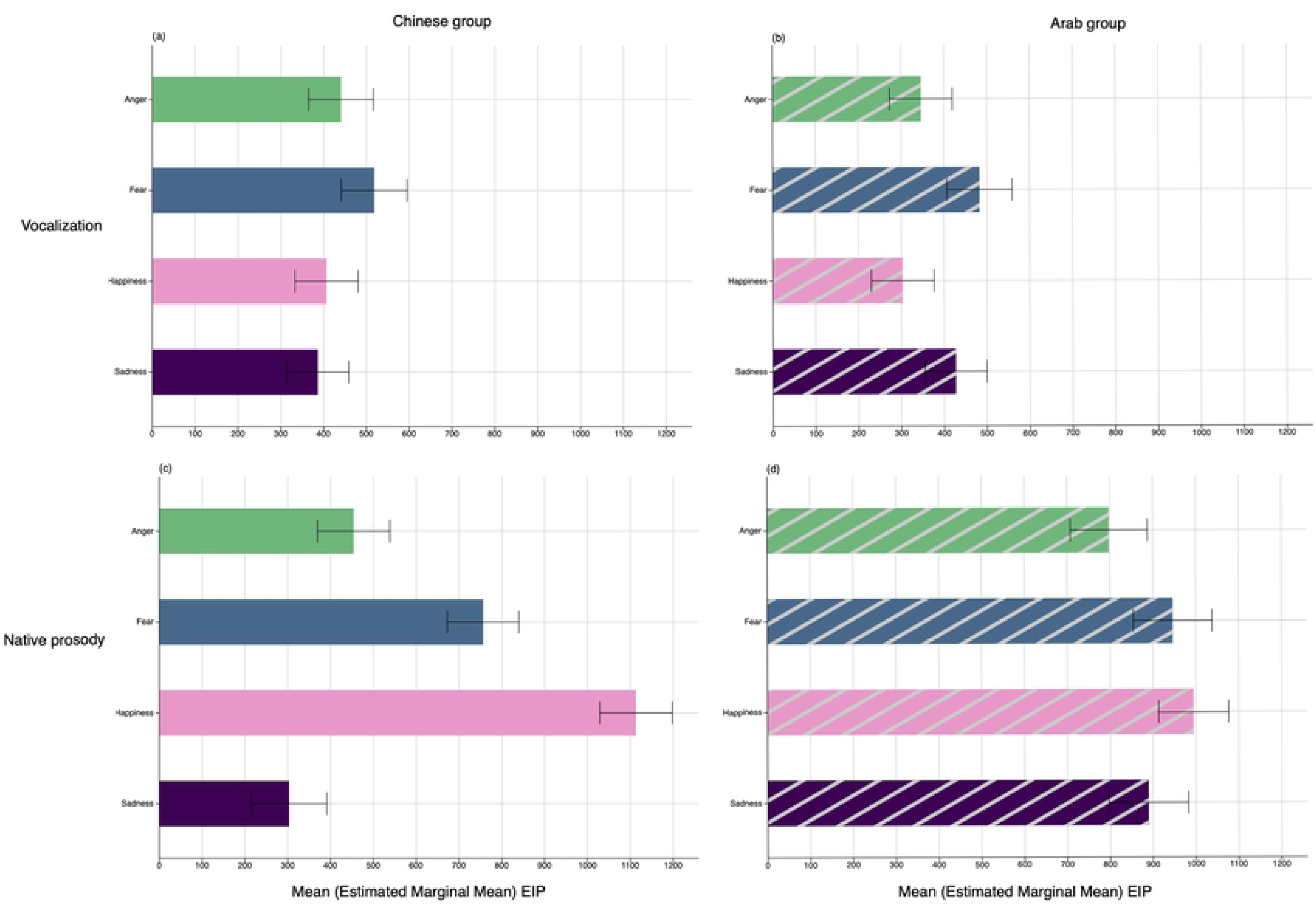
Recognition latency (EIP) by Event type and Emotion. Latency of each listener group to recognize emotions by vocal event type. The top panels show the Emotion Identification Points (estimated marginal means) when nonverbal vocalizations were presented and the bottom panels showing corresponding data when emotional prosody was presented in the listener’s native language (Mandarin-Chinese or Arabic). The performance of Chinese listeners is shown on the left and Arab listeners is shown on the right.

### How does emotion recognition unfold from native speech prosody?

#### Accuracy

Separate group analyses then considered the buildup of emotion representations when participants heard their native prosody (Chinese=Mandarin, Arab=Arabic; LMM: *HuScore (Native prosody) ∼ Emotion + Gate + Emotion*Gate + (1 | Subject*)). Accuracy differed by Emotion (Arab: *F* = 2.04, *df* = 3, *p* = 0.04; Chinese: *F* = 96.41, *df* = 3, *p* < 0.001), Gate duration (Arab: *F* = 89.04, *df* = 4, *p* < 0.001; Chinese: *F* = 51.08, *df* = 4, *p* < 0.001) and Emotion x Gate duration (Arab: *F* = 4.33, *df* = 12, *p* < 0.001; Chinese: *F* = 5.68, *df* = 12, *p* < 0.001; S4 Table a-b). Native prosody recognition improved incrementally as acoustic details accumulated but more gradually than for vocalizations, with significant improvements usually occurring at longer stimulus exposures (> 600ms, Fig. 1c-d). Moreover, the time course for recognizing specific emotions showed notable variability when Chinese vs. Arab listeners attended to acoustic features in their native language.

For the Chinese group, no significant gate-to-gate improvements occurred for any emotion until >600ms of the utterance was heard; between G600-GFull, recognition then increased significantly for certain emotions (fear, happiness, *p*s <.001). Chinese participants displayed superior recognition of sadness and anger from 200ms prosodic excerpts (anger = sadness > fear > happiness), a pattern that persisted up to 600ms. When Chinese listeners heard ungated utterances (GFull), all emotions were recognized at similar accuracy levels (Fig. 1c). For the Arab group, recognition of certain emotions (anger, happiness) improved when prosody increased from 200-400ms; otherwise, no significant gate-to-gate improvements were again observed until listeners heard >600ms of the utterance (G600-GFull), which was characterized by a sharp increase to detect *all* emotions (Fig. 1d). Contrary to Chinese listeners, Arab listeners displayed comparable accuracy for the four emotions at short stimulus exposures (G200-G600, except a slight “sadness” advantage at 200ms). Presentation of the full utterance produced marked improvements in recognition of Arabic prosody and differentiation of emotional meanings (happiness > anger = fear > sadness).

While these results show that prosodic representations build up gradually in each language and target hit rates did not tend to change when exposure times differed minimally (i.e., 100ms or 200ms), accuracy *did* improve significantly within each group when broader time windows were considered (e.g., when comparing G200-G600 or G400-GFULL, see post hoc contrasts in S4 Table a-b). Thus, prolonged analysis windows at later timepoints seemed to provide relevant acoustic details to recognize emotions from native prosody, in contrast to vocalizations which were isolated within a narrow time range early in the stimulus.

#### Latency

Direct group comparison of the EIPs for native prosody was then undertaken (LMM: *EIPtime (Native prosody) ∼ Group + Emotion + Group* Emotion + GFullDuration + (1 | Subject)*); S4 Table c). Recognition latencies depended on Group (*F* = 29.83, *df* = 1, *p*<.001), Emotion (*F* = 83.67, *df* = 3, *p* < .001), and Group x Emotion (*F* = 38.96, *df* = 3, *p*<.001). The time needed to isolate emotions from native prosody in the two languages spanned a sizable range (∼300-1100ms), averaging 600-1000ms of stimulus exposure. Overall, sadness (M = 597ms) and anger (M=626ms) were recognized faster than fear (M=852ms) and happiness (M=1054ms). These patterns varied somewhat in each listener group (Fig. 2), although happiness invariably required the most time to isolate (>1000ms). Anger, sadness, and fear were all recognized from significantly shorter excerpts in Mandarin than in Arabic (all *p*’s < .003), whereas happiness was recognized from shorter stimuli in Arabic vs. Mandarin (*p* = .049).

### Do listeners recognize emotions better from vocalizations than their native language?

Analyses then broadly examined whether vocalizations promoted a recognition advantage in accuracy or speed over native prosody through direct group comparisons (LMM for Accuracy: *HuScore (GFull) ∼ Group + EventType + Group*EventType + (1 | Subject)+ (1 | Emotion);* LMM for Latency: *EIPtime ∼ Group+ EventType + Group*EventType + GFullDuration + (1 | Subject)+ (1 | Emotion);* S5 Table a-b). The model performed on accuracy considered Hu scores at a single gate, GFull, when listeners had all acoustic information available to promote recognition of emotional targets.

#### Accuracy

Performance differed significantly by Event type (*F* = 25.95, *df* = 1, *p* < 0.001), pointing to superior accuracy to detect emotional vocalizations (M = 0.69) than native prosody (M=0.59) overall. A Group x Event type interaction (*F* = 9.90, *df* = 1, *p* < 0.001) qualified that while the group trends were similar (vocalization > native prosody), this pattern was significant for the Arab listeners (*p* <.001) but not the Chinese (*p* = .169, Fig. 3a). When listeners heard expressions in their entirety, there were no Group differences in the ability to recognize emotions in either the vocalization or the native prosody condition *(p*s > .191).

**Figure 3.**
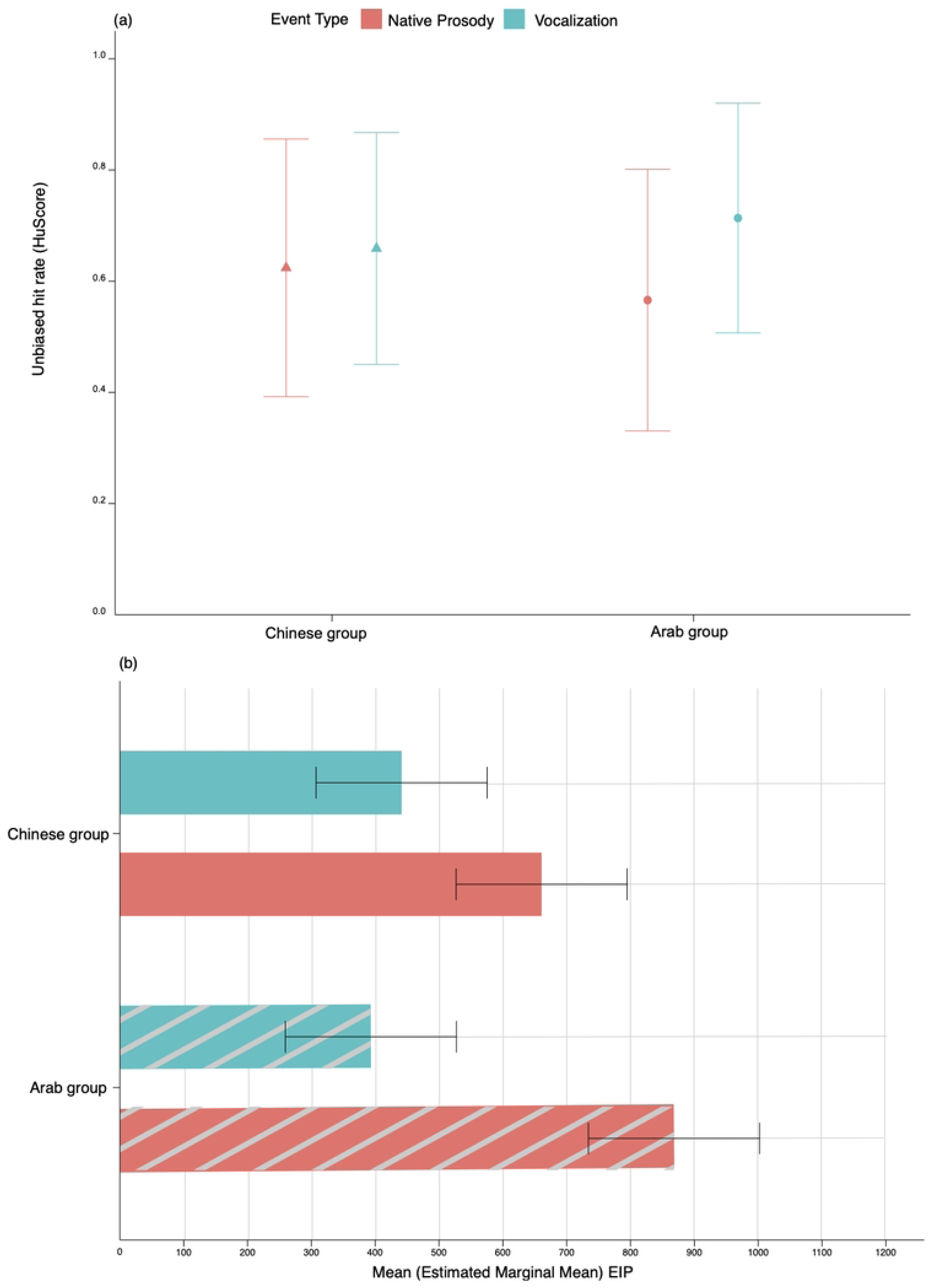
Recognition accuracy (HuScore) and Recognition latency (EIP) by Group and Event type. Recognition accuracy and latency across emotions by listener group and vocal event type. Fig. 3a (top panel) compares the accuracy (Hu score) of Chinese and Arab listeners to recognize vocalizations and native emotional prosody when expressions were presented in their entirety (GFull gating condition). Fig 3b (lower panel) shows the corresponding Emotion Identification Points (estimated marginal means) across emotions for the two listener groups and vocal event types.

#### Latency

The time needed to achieve stable recognition of emotions from vocalizations vs. native prosody varied by Group (*F* = 4.90, *df* =1, *p* = 0.03), vocal Event type (*F* = 489.77, *df* = 1. *p* < 0.001), and their interaction (*F* = 65.66, *df* = 1, *p* < 0.001). Overall, participants recognized vocalizations (M = 417ms) from significantly shorter acoustic excerpts than the same emotions expressed through native prosody (M = 765ms, *p*s < .001; Fig. 3b). Vocalizations were recognized at similar latencies by each group (*p* = .222), whereas native emotional prosody was identified from significantly shorter excerpts in Mandarin (Chinese listeners) than in Arabic (Arab listeners, *p <* .001, Fig. 3b).

### Does linguistic experience influence recognition of emotional prosody?

At a final step, separate group analyses considered patterns of emotion recognition in the three speech prosody conditions defined by the familiarity of each language to each listener group (native, L2-English, foreign), collapsed across emotion types (LMM: *HuScore ∼ Familiarity + Gate + Familiarity*Gate + (1 | Subject) + (1 | Emotion)*, S6 Table a-b). These models will reveal if representations underlying emotional prosody are facilitated by (native and/or second) language experience and *when* this knowledge comes into play during event processing^29^.

#### Accuracy

For Chinese listeners, accuracy depended on Gate duration (*F* = 147.3, *df* = 4, *p* < 0.001) and language Familiarity (*F* = 333.27, *df* = 2, *p* < 0.001) in the absence of an interaction. Emotional prosody recognition improved significantly within two principal time windows: between 200-400ms and when listeners had access to all acoustic information in the utterance (G600-GFull); these effects were observed consistently across speech contexts (native, L2, foreign) and were mirrored by the Arab group. Irrespective of stimulus exposure, Chinese listeners identified emotions more accurately from native prosody (Mandarin) than L2-English or foreign prosody (Arabic), exemplifying an ‘ingroup’ recognition advantage for native over non-native forms of prosody ^40^. Accuracy in the L2-English vs. foreign language conditions did not differ at any time point (Fig. 4a). Arab listeners displayed a unique pattern: accuracy depended on Gate duration (*F* = 186.25, *df* = 4, *p* < 0.001), language Familiarity (*F* = 52.77, *df* = 2, *p* < 0.001), and their interaction (*F* = 5.42, *df* = 8, *p* < 0.001). When prosodic stimuli were short (200-600ms), Arab listeners isolated emotions better in the *foreign* language, Mandarin, than from native (Arabic) or L2-English prosody, which did not differ. When the full utterance became available, Arab listeners showed marked improvements in the native and L2-English conditions, resulting in similar accuracy levels in the three speech prosody contexts.

**Figure 4.**
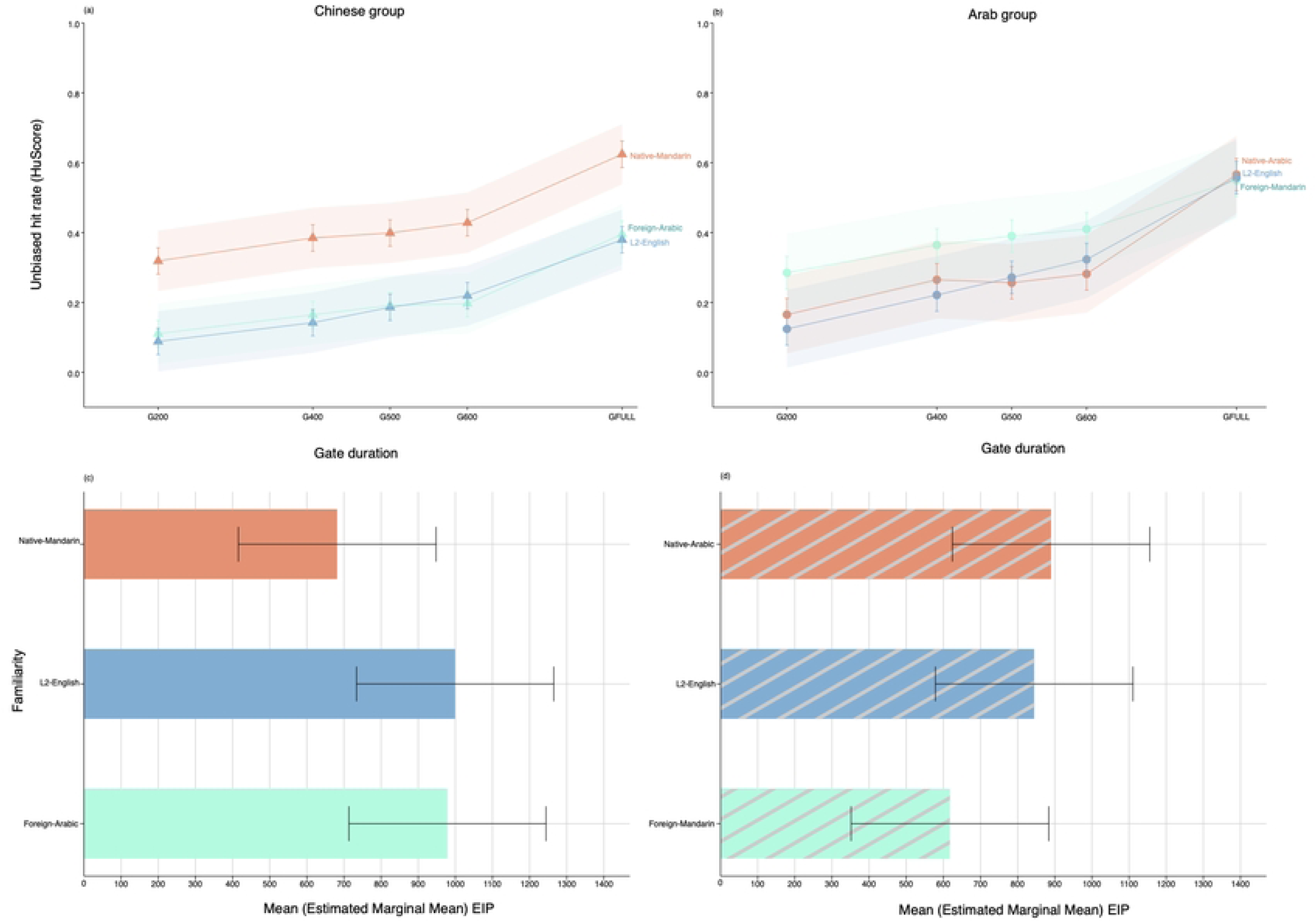
Recognition accuracy (HuScore) and Recognition latency (EIP) by Group and Speech familiarity. Effects of language familiarity on emotional prosody recognition as a function of stimulus exposure duration, across emotions by listener group. Fig. 4a (top panel) compares the accuracy (Hu score) of Chinese and Arab listeners to recognize emotional prosody over time in each group’s native language, L2-English (English as a second language), and in a foreign language. Fig 4b (lower panel) shows the corresponding Emotion Identification Points (estimated marginal means) across emotions for the two listener groups according to language familiarity.

To further contextualize these findings, we directly compared Group accuracy at GFull after recoding the speech prosody conditions according to the language of expression (Arabic, English, Mandarin) irrespective of its relevance/familiarity to a particular listener group (LMM: *HuScore (GFull) ∼ Language + Group + Language*Group + (1 | Subject) + (1 | Emotion),* S6 Table c). Recognition of full prosodic expressions did not differ overall by Language, but differed by Group (*F* = 5.59, *df* = 1, *p* = 0.02) and Group x Language (*F* = 28.73, *df* = 2, *p* < 0.001). Chinese listeners displayed significantly better recognition of Mandarin emotions from full utterances (Mandarin > English = Arabic), whereas Arab listeners displayed no advantage across languages for full utterances (Mandarin = English = Arabic). Emotional prosody in Arabic and English were recognized more accurately by Arab than Chinese listeners, whereas the opposite was true for Mandarin (Chinese > Arab, Fig. 4a).

#### Latency

Group comparison of EIPs in the three prosody conditions (LMM: *EIPtime ∼ Group + Familiarity + Group*Familiarity + GFullDuration + (1 | Subject) + (1 | Emotion)*; S6 Table d) uncovered effects of Group (*F* = 9.56, *df* = 1, *p* = 0.003), language Familiarity (*F* =24.71, *df* = 2, p < 0.001), and Group x Familiarity (*F* = 103.29, *df* = 2, *p* < 0.001). Chinese listeners had significantly shorter recognition latencies for native prosody (M = 682ms) than foreign-Arabic or L2-English prosody (M = 979ms and 999ms, respectively, *p*s < .001). In contrast, Arab listeners demonstrated earlier recognition points for foreign-Mandarin prosody (M = 618ms) than native-Arabic and L2-English (M = 890ms and 845ms, respectively, *p*s < .001). Thus, emotion representations always stabilized more quickly when listening to speakers of Mandarin irrespective of familiarity (Fig. 4b). As noted earlier, Chinese listeners displayed faster recognition of native prosody (*p* <.001), but Arab listeners had shorter EIPs than Chinese listeners when judging foreign prosody and L2-English (*p’s* <.001).

## Discussion

Research suggests that auditory expressions of emotion are recognized more “successfully” as vocalizations—i.e., through grunts, sighs, laughs, sobs, and other brief non-linguistic sounds—than through speech prosody. As phylogenetically older and more reflexive signals ^20^, spontaneous vocalizations correlate with autonomic and physiological changes in the speaker’s internal state and are produced with less cognitive control and with less acoustic constraints than speech ^14,18,51^. These factors seem to boost the attentional salience of nonverbal expressions ^3,52^ and promote the perceptual “clarity” of discrete emotions encoded by nonverbal signals over speech-embedded emotion expressions ^2,4,9^, although there is still a paucity of work that directly compares recognition of the two event types.

Our data provide within-subjects verification that listeners are more accurate and require substantially less time to detect basic emotions communicated by vocalizations than prosodic features of their native language. When exposed to ungated exemplars of each event type, vocalizations were recognized at a superior level to native speech overall (0.69 vs. 0.59, respectively). Even more notable, listeners required nearly *half* the exposure time on average to form a stable impression of the speaker’s emotion from vocalizations (Mean =417ms) than from native prosody (Mean = 765ms). Thus, it can be said that listeners in our study achieved an accurate sense of the intended meaning conveyed by vocalizations more *efficiently*, i.e., in a much shorter time period, than for speech prosody. In evolutionary terms, it is thought that vocalizations were functionally purposed to communicate rapid and ‘honest’ details to conspecifics about emotional events necessary for survival which require minimal conceptual elaboration ^53^. Our findings also suggest that the primary communicative function and main ecological benefit of vocalizations is to transmit acoustic information that allows listeners to automatically construct a fine-grained, categorical representation of the speaker’s emotion state in an expedited manner ^14,54,55^. Reducing the latency of emotion recognition would accelerate adaptive action tendencies associated with discrete emotional signals ^1^, especially when visual cues are impoverished, and allow earlier and clearer *predictions* about the emotional situation.

Anger vocalizations (e.g., shouts) displayed an early detectability advantage and were almost fully recognized from 200ms bursts ^38^. Increasing stimulus exposure to 400ms then resulted in near peak accuracy for all vocalization types (with the exception of sadness/crying which improved somewhat beyond 400ms). These patterns underscore that structural differences within the first 200-400ms of vocalizations (beginning <200ms) provide adequate perceptual information to discriminate and form abstract categorical representations of the emotion qualities communicated by nonverbal acoustic signals ^54,55^. Calculation of EIPs confirmed that the functional significance of all vocalizations was firmly established within a narrow ∼300-500ms time window ^36^ for both Chinese and Arab listeners, who displayed minimal performance differences (despite the fact that vocalizations were extracted from different recording databases, produced by speakers of various European languages). Arab/Chinese listeners only differed in their recognition latencies for happiness/laughter, although isolation points for this emotion were still rapid (300-400ms). These data bolster claims that nonverbal vocalizations possess robust ‘universal’ acoustic elements ^57^ that reveal their functional significance to perceivers 300-500ms post-onset of the vocalization. This process can be achieved without prior experience and shows little cultural variation, except possibly for positive emotions ^8,9^.

### Effects of language on emotional prosody recognition

Emotional prosody was marked by group differences in accuracy, timing, and unique recognition trajectories in each native language, pointing to a more pronounced impact of linguistic and socio-cultural variables on how emotional representations are constructed in speech. When speech excerpts were limited (200-600ms), Chinese listeners displayed superior detection of anger and sadness from native prosody, while Arab listeners displayed a slight advantage for sadness ^12,29,34^. The Arab group was notably less accurate to recognize emotions from native prosody when stimulus exposure was short (<600ms). Prosody recognition significantly increased and stabilized only when listeners were fully exposed to native pseudo-utterances (G600-GFull), yielding moderately high emotion hit rates in each group that are comparable to the literature (∼ 3x chance accuracy level, e.g., ^7,10^). Chinese listeners recognized all emotions with similar accuracy when all acoustic details were available, whereas Arab listeners were more accurate for certain emotions (happiness > anger = fear > sadness). The timing (EIP) data show that emotions were isolated from prosody at vastly different rates, ranging from 300-1100ms in Mandarin compared to 800-1000ms in Arabic (EIPs in each language were always shortest for sadness and anger and longest for happiness, cf. ^29,32,33^). An important conclusion that can be drawn from these data is that emotional prosody recognition is interdependent on language context and structure ^10^, and that refining impressions of a speaker’s emotional state from speech often benefits from phrase-final acoustic information ^33–35^. In contrast, we found little evidence that event details beyond 600ms post-onset of emotional targets aided recognition of vocalizations for our paradigm.

Our data highlight two critical time periods which may be crucial for extracting details about vocal emotion expressions following event onset: an early window (∼0-400ms), which promotes full recognition of nonverbal vocalizations and appears to allow rough differentiation of emotionality/highly salient emotional qualities communicated by speech prosody ^3,15,24,36^; and a late extended window (400-1000ms+), which is necessary to monitor and consolidate cues that encode the emotional meaning of prosodic expressions as speech unfolds ^29,31,32,34^. The late integration window also serves to incorporate semantic information relevant to the speaker’s emotion state ^56,57^ and considers phrase-final acoustic cues which impact on how emotional prosody is contextually interpreted ^7,34,35^. This processing scheme ensures rapid bottom-up detection of the categorical relevance of motivationally salient vocal signals from acoustic-perceptual information in the early processing interval (including trait impressions of dominance, attractiveness, etc. ^58^). At the same time, it explains the *gradual* emergence of stable prosodic representations in speech which tended to build up over longer sampling periods in our data. On average, the EIP data show that the emotional significance of speech prosody—at least for basic negative emotions such as anger, fear and sadness—is reliably established ∼500-800ms post-onset of an utterance ^29,31–33^. However, unlike vocalizations, these estimates vary considerably across items and languages and can be substantially longer when more socially-constructed emotions, including happiness, are studied (EIPs >1-2 seconds) ^32,34^.

Theories of speech perception have proposed that auditory-perceptual integration of basic linguistic units (segments vs. syllables) is accomplished by distinct brain mechanisms that sample acoustic information over different time scales (25-80ms windows for segments, 150-300ms windows for syllables ^59^). Along these lines, different forms of vocal signals (vocalizations, prosody) may reveal emotional meanings at unique timepoints owing to distinctly adapted procedures for sampling emotionally relevant acoustic variation over different time scales. Arguably, recognizing emotions in speech depends on a broader analysis window that roughly aligns with major perceptual units that promote linguistic comprehension, such as syllabic units ^5,32^. As listeners appraise emotional speech cues, they must also distribute perceptual and cognitive resources to extract linguistic meaning from the acoustic signal; the conscious nature and complexity of this process may contribute to why listeners build more gradual representations of the speaker’s emotion state from prosody when compared to vocalizations. Humans also learn that the *antecedents* of emotional events that are typically expressed in fluent speech are rarely as urgent as those signalled by vocalizations (see ^60^ for a discussion), and that speakers often intentionally manipulate emotional cues to serve their interpersonal goals in the communicative situation ^53,61^. Acquired knowledge of these form-function relationships could explain the more variable and sustained analysis period associated with emotional prosody recognition, as it is believed that the human capacity to communicate using language, and to communicate *emotions in language,* co-evolved ^62^.

### On the ingroup advantage for emotional prosody recognition

Several variables are likely to shape how speech-embedded emotions, as well as volitional productions of vocalizations such as laughter ^63,64^, are assigned value over time and “recognized” in daily interactions; these include cultural preferences/display rules and inferences about the strategic “pragmatic” goals of the speaker in the context of emotional communication ^53,65^. Of main interest here, there is mounting evidence that language experience familiarizes people with emotional dialects or vocal expressive “styles” used by particular groups ^39^, allowing more precise and more rapid construction of emotional prosody representations in listeners’ native vs. a foreign language (ingroup advantage ^40^). Our study provides a unique view on this issue, as we employed a fully crossed presentation design involving participants who had no familiarity with the foreign language (Mandarin vs. Arabic) but who all had shared proficiency in English as a second language acquired from childhood, allowing graded effects of familiarity to be evaluated in each group.

Our findings replicated the ingroup advantage for only one of our two groups: Chinese listeners were more accurate and required less time to name emotions in Mandarin than the other two languages (native > foreign = L2-English, ^7,41,66^). This relationship was established after hearing only 200ms speech excerpts and did not temporally evolve (cf. ^29^). Presumably, Chinese listeners isolated emotional meanings more efficiently in Mandarin because these stimuli abided by culturally acquired norms of expression familiar to the participants; since performance in L2-English and Arabic (foreign) did not differ, this effect is unlikely to be traced to basic problems in phonological encoding for non-native stimuli.

There was no evidence that Arab participants presented an ingroup advantage in accuracy or speed at any stimulus duration (see ^67^ for similar conclusions using a fully crossed design). Moreover, unlike the Chinese performance of the Arab group changed as a function of acoustic exposure in each language. When excerpts were brief (200-600ms), Arab listeners were unexpectedly superior in Mandarin (foreign > native = L2-English) and EIPs were consequently shorter in Mandarin than the other languages overall. However, when Arab listeners judged full utterances emotion accuracy was comparable in the three languages (Arabic = Mandarin = L2-English). These patterns show that they relied heavily on utterance-final acoustic cues to form stable impressions of emotion in Arabic and L2-English, and ultimately, Arab listeners achieved higher recognition rates in these two language contexts than Chinese listeners. However, their *path* to recognition and how they dynamically integrated emotion-related cues in each language was distinct.

Counter-intuitively, our results point out that emotional prosody may be recognized more efficiently at times in a completely unfamiliar language, overriding any advantages conferred by experience with emotional dialects used by native speakers. Although we strived to match our prosodic recordings in the three languages along key dimensions, it seems clear that Mandarin speakers supplied more distinct and representative cues to emotion *at earlier stages* in their productions than the Arabic or English speakers, allowing recognition to stabilize more quickly in this language context for *both* native (Chinese) and foreign (Arab) listeners, irrespective of familiarity. However, given the sustained time course of emotional prosody analysis, the initial (bottom-up) advantage to recognize Mandarin prosody was eliminated for Arab listeners when acoustic information became available towards the end of Arabic/English utterances. These patterns remind us that the task of communicating emotions both within and across linguistic boundaries depends not only on top-down factors, i.e., acquired cultural knowledge and contextual expectations about a speaker’s vocal behaviour; it is fundamentally driven by the *dynamic* quality of the input, i.e., how well speakers execute universally shared ‘affect programmes’ ^68^ that guide emotional speech recognition at different timepoints, and how well acoustic cues facilitate emotional “inference rules” shared by the speaker-listener ^10^. In daily life, speakers encode emotion with different levels of intensity ^69,70^ and not all individuals are adept at vocally encoding recognizable emotion states in speech ^5^. As this research moves forward, the interplay of stimulus-related features and various knowledge sources that are brought to bear on the act of emotional speech recognition will come increasingly to light.

Our findings supply no clear evidence that knowing a second language benefits emotional prosody recognition, despite the high English proficiency of our participants. Performance in the L2-English condition mirrored the pattern of foreign prosody (Chinese group) or native prosody (Arab group; cf. ^29^ for data on English and Hindi). Bhatara et al. ^42^ reported that L2-English proficiency in a group of French listeners had no effect on prosody recognition (for negative emotions) or seemed to *interfere* in this process (for positive emotions). Other studies suggest that emotional prosody recognition from single words is equally accurate and rapid in one’s native language and L2-English ^43^. Our data call for continued monitoring of the relationship between L2 proficiency, emotion recognition, and how acquiring “prosodic-pragmatic competence” in a second language is influenced by different forms of instructional practice ^71^.

## Conclusions

In closing, it should be emphasized that the recognition of emotions from vocalizations is *not* necessarily more accurate or precise than for speech prosody, if the quality *and* quantity of input listeners receive from each form of expression are adequate. Here, Chinese listeners were just as accurate to identify basic emotions from vocalizations and from native prosody when acoustic exposure to each event was unrestricted; thus, one can say with confidence that a subset of basic emotions can be communicated with equal “success” (i.e., accuracy) through either nonverbal or linguistic vocal channels. However, emotional meanings were always understood much more quickly from nonverbal sounds than prosody ^3^. Ensuring that robust representations of a speaker’s emotion state are arrived at *efficiently* may be the core function of vocalizations that distinguishes the two vocal communication subsystems.

## Acknowledgements

This research was supported by Insight Grant #435-2022-0391 from the Social Sciences and Humanities Research Council of Canada to Marc D. Pell. We thank Jamie Russell for her valuable help in stimulus preparation and pilot testing.

